# Single-cell RNA-seq reveals intrinsic and extrinsic regulatory heterogeneity in yeast responding to stress

**DOI:** 10.1101/179093

**Authors:** Audrey P. Gasch, Feiqiao Brian Yu, James Hose, Leah E. Escalante, Mike Place, Rhonda Bacher, Jad Kanbar, Doina Ciobanu, Laura Sandor, Igor V. Grigoriev, Christina Kendziorski, Stephen R. Quake, Megan N. McClean

## Abstract

From bacteria to humans, individual cells within isogenic populations can show significant variation in stress tolerance, but the nature of this heterogeneity is not clear. To investigate this, we used single-cell RNA sequencing to quantify transcript heterogeneity in single *S. cerevisiae* cells treated with and without salt stress, to explore population variation and identify cellular covariates that influence the stress-responsive transcriptome. Leveraging the extensive knowledge of yeast transcriptional regulation, we uncovered significant regulatory variation in individual yeast cells, both before and after stress. We also discovered that a subset of cells decouple expression of ribosomal protein genes from the environmental stress response, in a manner partly correlated with the cell cycle but unrelated to the yeast ultradian metabolic cycle. Live-cell imaging of cells expressing pairs of fluorescent regulators, including the transcription factor Msn2 with Dot6, Sfp1, or MAP kinase Hog1, revealed both coordinated and decoupled nucleocytoplasmic shuttling. Together with transcriptomic analysis, our results reveal that cells maintain a cellular filter against decoupled bursts of transcription-factor activation but mount a stress response upon coordinated regulation, even in a subset of unstressed cells.

## INTRODUCTION

When adversity strikes, it is often the case that some cells in an isogenic population survive whereas others do not. Such phenotypic heterogeneity has been observed in isogenic microbes exposed to environmental stress as well as normal and malignant human cells surviving chemotherapy drugs (1-7). While genetic mutations can produce cells with heritably high stress tolerance, in many cases the heterogeneity is transiently induced by epigenetic processes (8, 9). For example, some isogenic cells within cultures of *Saccharomyces cerevisiae* can survive extreme heat stress whereas most cells in the culture cannot (10). Stress-tolerant individuals may be in an altered state, since they often display transiently reduced growth and markers of the stress response (1, 10-12). But whether this state mimics that of stress-treated cells that have fully mounted a stress response or instead emerges from partial, stochastic events is unclear. Understanding what gives rise to cell-to-cell heterogeneity in stress survival has broad application, from treating pathogenic microbial infections to blocking drug-resistant human metastases.

In several systems, variation in stress tolerance can be traced to heterogeneous expression of defense genes. Graded expression of the stress-responsive *TSL1* gene in unstressed yeast quantitatively predicts how well individual cells in a culture will survive severe heat (10). In some cases, such variation is ‘intrinsic’ to the gene promoter: many defense genes are transcribed through TATA-dependent promoters (13, 14), which produce stochastic transcriptional bursts proposed to play a role in bet hedging (15-19). But ‘extrinsic’ variation in the cellular system, *e.g.* activation of the broader upstream signaling response or transition through other physiological states, likely plays an important role. Stewart-Ornstein *et al.* showed that targets of the yeast ‘general-stress’ responsive transcription factor Msn2 often behave coordinately in single cells, suggesting concerted activation of the entire regulon even in the absence of added stress (20). Targets of several other transcription factors also showed coordinate behavior across single cells, suggesting that the variation may emerge from stochastic activation of Protein Kinase A (PKA), a common upstream regulator of those factors (20).

In response to acute stress, Msn2 is activated as part of a broader signaling network that regulates the Environmental Stress Response (ESR), a common transcriptomic response triggered by diverse stresses (21, 22). The ESR includes induced expression of ∼300 defense genes, regulated in part by Msn2 and its paralog Msn4, which is coordinated with repression of ∼600 genes encoding ribosomal proteins (RPs) and factors involved in ribosome biogenesis and other processes (RiBi genes). RP and RiBi genes are highly transcribed in actively growing cells, but repressed in response to stress through release of the RP activator Sfp1 or recruitment of the RiBi transcriptional repressors Dot6/Tod6 and histone deacetylases (23-27). Activation of the ESR after mild stress can impart increased tolerance to subsequent stress, known as acquired stress resistance (28-30). In some cases, the ESR program also correlates with reduced growth rate, most notably in nutrient-restricted chemostats and in slow-growing mutants potentially experiencing internal stress (31-35). In fact, O’Duibhir proposed that the ESR may simply be a byproduct of cell cycle phase, since slow-growing mutants with prolonged G1 display an ESR-like transcriptome profile (35). In nearly all studies to date, increased expression of the induced-ESR (iESR) genes is coupled to reduced expression of RP and RiBi genes in the repressed-ESR (rESR) gene set. Whether regulation of the iESR and rESR genes can be decoupled in wild-type cells is unclear.

Msn2 over-expression is sufficient to induce multi-stress tolerance in yeast cells (36). Thus, cell-to-cell variation in Msn2 activation could explain the heterogeneity in stress tolerance in an actively growing culture. But it remains unclear if this variation correlates with broader transcriptome changes, if the magnitude of the response in unstressed cells mimics that seen in stressed cells, or if fluctuations in the response correlate with cell-cycle phase. Here, we addressed these questions through single-cell RNA-sequencing (scRNA-seq) coupled with single-cell profiling of transcription factor activation dynamics, before and after stress. Our results reveal variable activation of both the ESR and condition-specific stress regulators after stress, and heterogeneity in ESR activation before stress due to both coordinated and discordant induction of ESR regulators. While ESR activation shows no relation to cell-cycle phase in unstressed cultures, we found that some cells decouple regulation of RP transcripts in a manner linked to S-phase but apparently unrelated to expression changes associated with the yeast metabolic cycle.

## RESULTS

We used the Fluidigm C1 system to perform scRNA-seq on actively growing yeast cells collected from rich medium, before and 30 min after treatment with 0.7 M sodium chloride (NaCl) as a model stressor. Although the Fluidigm system generally profiles fewer cells than other methods, it has a substantially higher capture rate enabling deeper investigation of the cellular transcriptome (37, 38). Cells were immediately fixed by flash freezing to preserve the transcriptome during the capture process. We optimized a protocol to capture partially spheroplasted yeast cells on the C1 platform, ensuring that cells remained intact during capture but lysed in the instrument. We performed two C1 chip runs for each of the unstressed and stressed cultures and selected 85 and 81 captured single-cells respectively; 83 and 80 yielded successful libraries (see Methods). Libraries were pooled, paired-end fragments were sequenced, and identical fragments collapsed to a single count to minimize amplification biases, producing a median of 1.4 million mapped de-duplicated fragments per cell (see Methods). Each transcriptome encompassed 735-5,437 mRNAs (median = 2,351), with a total of 5,667 out of the 5,888 yeast transcripts (39) covered by at least 5 reads in ≥5% of cells (Table 1). As is well known for scRNA-seq, low abundance transcripts with fewer read counts displayed a lower detection rate (*i.e.* lower fraction of cells in which the transcript was measured), likely a result of both technical noise and true biological variation (38, 40-42). The averaged responses of all stressed cells compared to all unstressed cells agreed well with bulk measurements (correlation between log_2_(fold change) = 0.7), especially for stress-regulated genes (see Fig S4), validating our procedure. For much of our analysis, we focused on the log_2_(normalized read counts) for each transcript in each cell, scaled to the mean log_2_(normalized read counts) of that mRNA in all other cells in the analysis (referred to as ‘mean-centered log_2_(read counts)’ or ‘relative log_2_ abundance’). Thus, positive log_2_ values indicate expression above the population mean of that transcript, and negative values represent expression below the mean.

### Quantitative variation in ESR expression

As expected, stressed and unstressed cells could be readily distinguished based on their cellular transcriptome, primarily driven by expression of the ESR genes (Fig 1A). Most unstressed cells displayed relatively high abundance of RP and RiBi transcripts and low abundance of iESR transcripts, consistent with ESR suppression, whereas stressed cells displayed the opposite patterns indicative of ESR activation. However, there was considerable variation in the magnitude of ESR activation, both before and after stress. Some stressed cells showed concertedly stronger activation of the ESR than other cells (Fig 1B-C). Among unstressed cells, at least 4% showed mild ESR activation, as evidenced by low RP expression relative to other unstressed cells (FDR <0.05, T-test, see Methods) coupled with high relative iESR mRNA abundances (Fig 1C, asterisks). In general, quantitative differences in ESR activation were correlated across ESR subgroups: cells with higher relative iESR transcript abundance generally showed lower relative RiBi and RP mRNA levels, whereas quantitative differences in RP abundance were generally correlated with RiBi abundances, especially in stressed cells (Fig 1B). The quantitative and correlated variation across these groups suggests coordinated cell-to-cell variation in ESR activation levels, both before and after stress.

**Figure 1.**
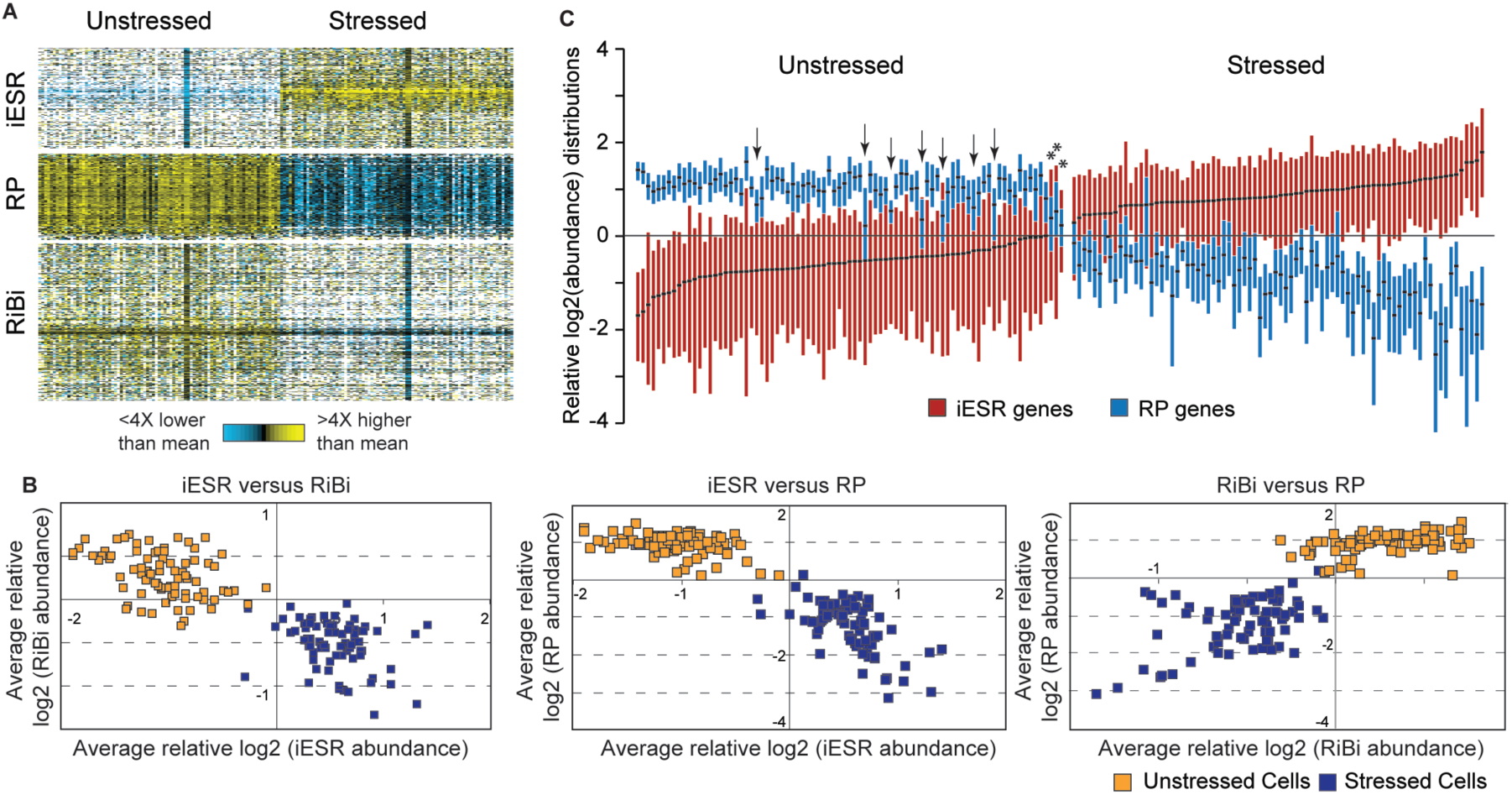
Quantitative variation in ESR activation across cells. (A) Mean-centered log_2_(read counts) for ESR gene groups before and after stress. Each row represents a transcript and each column is an individual cell, with expression values according to the key; white indicates no detected transcript. (B) The average mean-centered log_2_ values for a given ESR gene group as measured in one cell was plotted against the average mean-centered log_2_ values for a second ESR gene group as measured in the same cell. (C) Boxplots (without whiskers) of mean-centered log_2_(read counts) of RP and iESR transcripts in individual cells, sorted by iESR-group median. Arrows indicate unstressed cells with unusually low RP transcript abundances (FDR < 0.05, see text) and asterisks indicate those cells that also had high median iESR log_2_ values.

But there was also evidence of group-specific variation that appeared decoupled from ESR activation. Of the twelve percent of unstressed cells with lower relative RP abundance (FDR < 0.05, T-test), two thirds did not show significant iESR activation or RiBi repression (Fig 1C arrows and Fig S1). Likewise, a subset of stressed cells with strong iESR induction had significantly weaker repression (*i.e.* higher relative abundance) of RP transcripts (FDR < 0.05, T-test, see more below). These results suggest that heterogeneity in other cellular co-variates may influence expression of RP transcripts separate from full ESR activation, investigated in more detail below.

### Low variation in RP transcripts reflects tight cellular control

In the process of this analysis, we noticed that RP mRNAs showed tight distributions with relatively low variance in unstressed cells (Fig 1B and 1C). In fact, RPs showed among the lowest variances across the range of transcript abundances, whereas RiBi and iESR transcripts did not (Fig 2). The distinction persisted in stressed cells, although RP variance was notably higher after stress treatment (Fig 2C, D). We also noticed that the detection rate for RP transcripts appeared to be exceedingly high, even for RP transcripts known to be expressed at low abundance (43). To investigate this, we plotted the detection rate versus read counts per transcript length as a proxy for mRNA abundance and devised a statistical test based on cubic splines to identify differences across gene groups (see Methods). As a group, RP transcripts (p < 0.0028), and to a lesser extent RiBi mRNAs (p< 0.025), showed significantly higher detection rates compared to randomly chosen genes, for both stressed and unstressed cells (Fig 3). The result was not an artifact of known covariates of RP mRNAs. Although RP mRNAs are generally short, they remained statistically different from randomly chosen transcripts that are shorter than the median RP length (p<0.0012). Many RPs also have close paralogs in the genome, which could obscure true abundance if reads mapping to multiple locations are discounted from the alignment. But RPs remained significant, at least for unstressed cells (p = 0.02), when the analysis was performed only on mRNAs without a close paralog (BLAST E value > 1e-5). We validated the difference in detection rate for several transcripts using single-molecule mRNA fluorescence *in situ* hybridization (FISH, Fig 4). *RLP7*, encoding a ribosome-associated RP-like protein, and phosphatase-encoding *PPT1* mRNA were sequenced to similar read densities, but were detected in 87% versus 53% of unstressed cells, respectively. The distinction was confirmed by FISH: *RLP7* was measured in 78% and 50% of cells collected before and after stress, whereas *PPT1* was measured in 50% and 30% of cells, respectively, despite similar abundance ranges when present.

**Figure 3.**
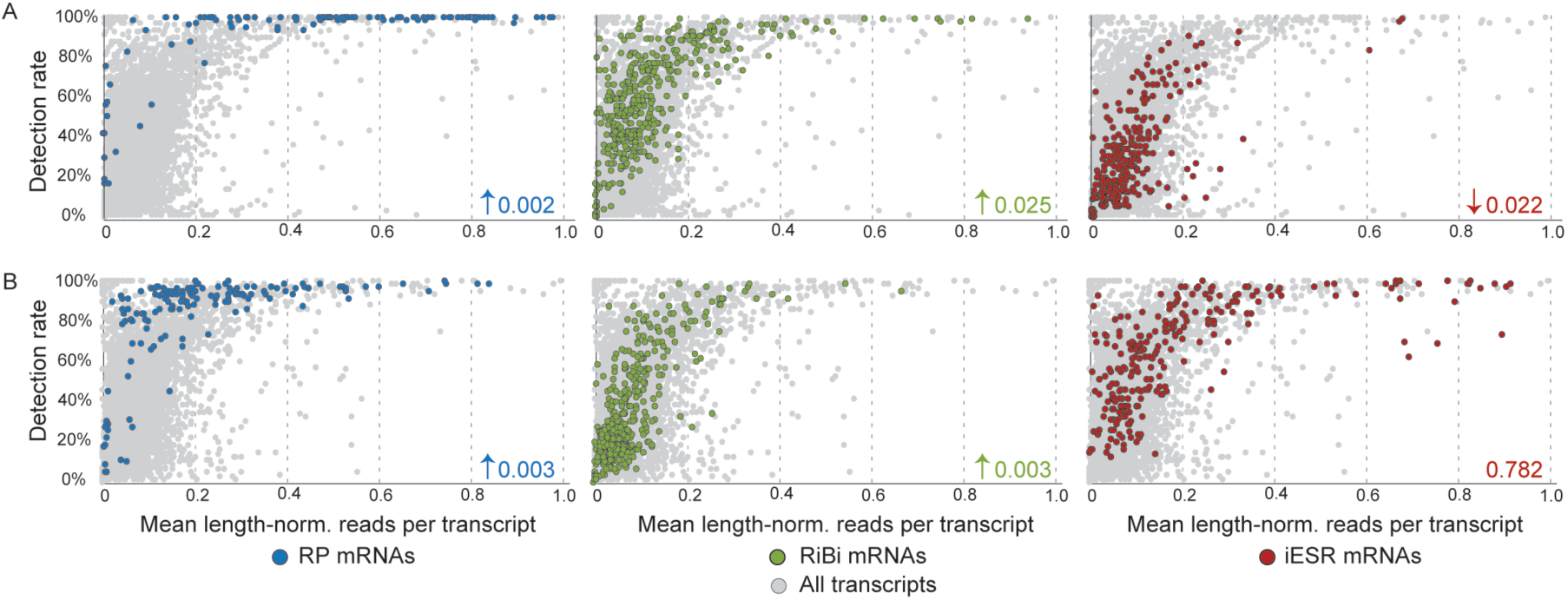
Transcript detection rate correlates with functional class. The fraction of cells in which each mRNA was detected was plotted against the mean length-normalized read count for that transcript, calculated from cells in which the transcript was measured, in (A) unstressed or (B) stressed cells. Listed p-values and arrows indicate if the detection rate was higher or lower than randomly sampled genes. Plots are zoomed in to show transcripts in the lower-read range; most transcripts above this range are measured in all cells, not shown.

**Figure 4.**
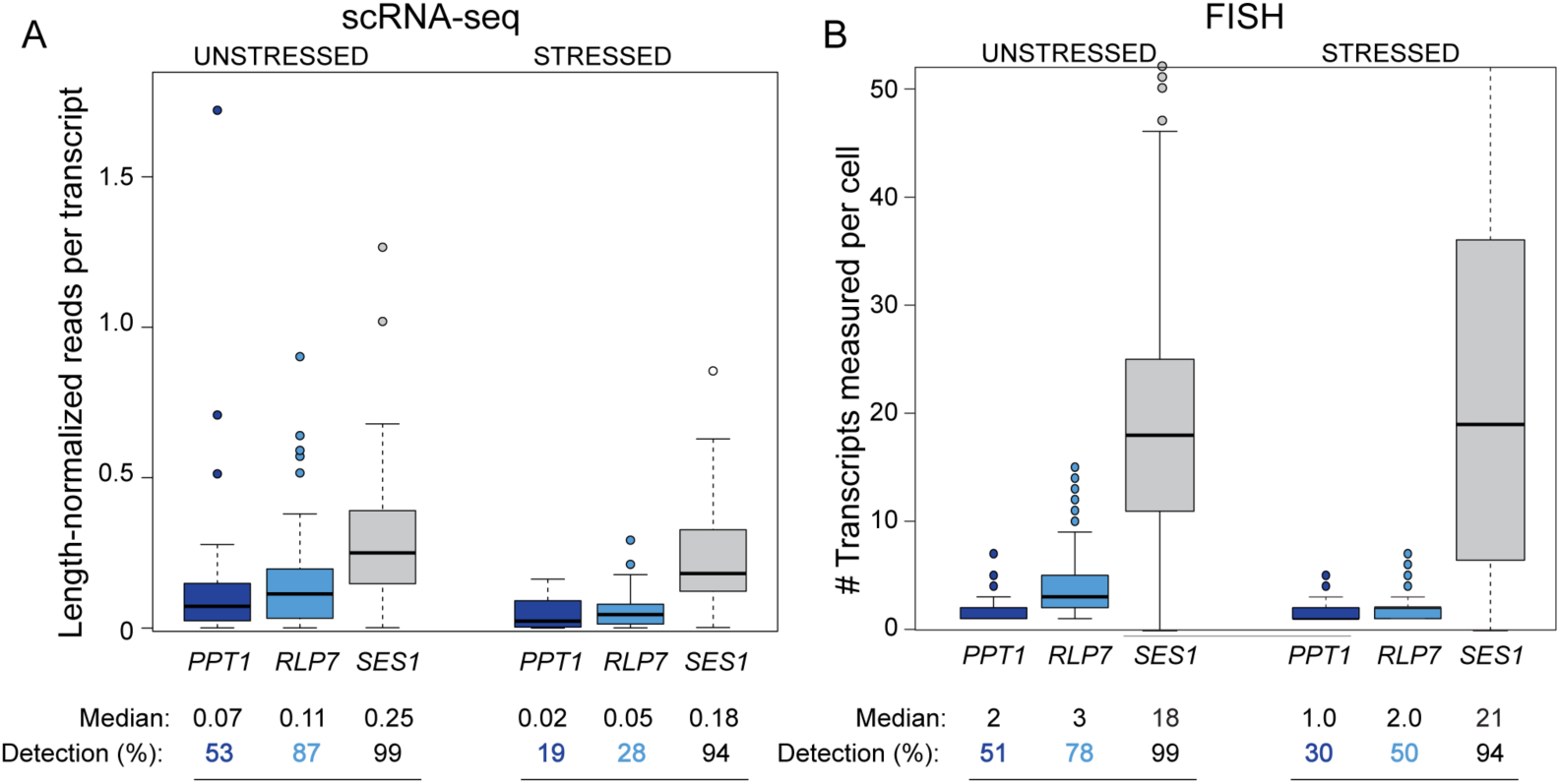
Single-molecule FISH confirms differences in detection rate at several transcripts. Distributions of (A) length-normalized read counts measured by scRNA-seq and (B) mRNA molecules per cell measured by single-molecule FISH, for *PPT1*, *RLP7*, and *SES1* as a control. Note only part of the *SES1* distribution is shown. Median counts in cells with a measurement and detection rate (% of cells with a measurement) are listed below the figure.

**Figure 2.**
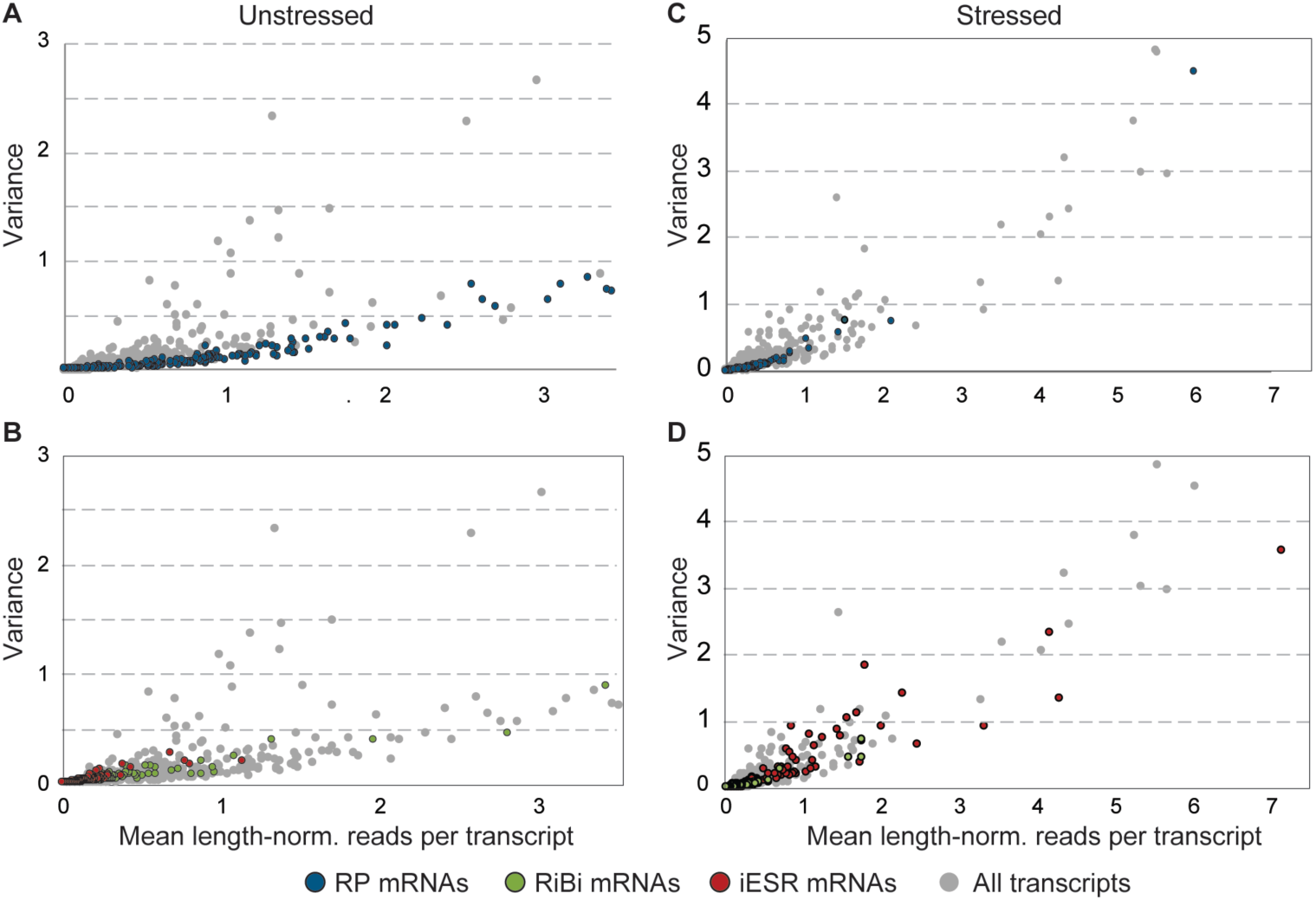
RP transcripts show low variation in abundance across cells. The mean and variance of transcript read counts per mRNA length (‘length-norm’) was plotted for each mRNA from unstressed (left) or stressed (right) cells. (A,C) highlight RP transcripts and (B,D) highlight iESR and RiBi transcript against all other mRNAs (grey points). Plots are zoomed to capture most points.

One likely reason for the tight control on RP transcripts is the importance of stoichiometric protein expression for proper ribosome assembly, and several RP transcripts are subject to extensive regulation to impose this control (44-47). We sought other transcripts whose detection rate was higher than predicted by their abundance and identified mRNAs (lacking paralogs) that were above the RP-fit spline (Table 2). Remarkably, this group was heavily enriched for mRNAs encoding multi-subunit protein complexes (48), in the analysis of both unstressed (p = 3.5e-4) and stressed cells (p = 2e-25, hypergeometric test). This included subunits of the proteasome, chaperonin-containing T complex, nuclear pore, as well as mRNAs encoding proteins destined for various subcellular regions. We also found mRNAs at the opposite end of the spectrum: iESR mRNAs displayed lower-than-expected detection rates, before – but not after – stress (p = 0.022). Many iESR genes are regulated by burst-prone TATA-containing promoters (13, 15-17), and indeed other TATA-regulated genes without paralogs were weakly enriched among those with unusually low detection rates (p = 0.01). Together, these results confirm that the detection rate is not merely a function of technical variation (38, 40) and show that transcripts in different functional groups are subject to different regulatory constraints per cell.

### ESR activation does not fluctuate with cell cycle phase

The analysis in Fig 1 revealed variation in ESR activation across cells, both before and after stress, as well as decoupling of RP expression in some cells. We hypothesized that this variation could emerge if cells are in different physiological states. The first candidate was cell-cycle phase. We used the program Pagoda (49) to identify clusters strongly enriched for known cell-cycle mRNAs, and then classified cells based on the expression peaks of the cluster centroids (Fig 5A, Table 3, see Methods). A subset of cells could not be classified, in part because we were unable to identify coherent expression among M-phase genes. Comparing the fraction of cells in each phase before and after stress recapitulated the known G1 delay after osmotic stress (50, 51)(Fig 5B). Interestingly, many more cells could not be classified based on their expression profile after NaCl treatment; while some of these cells could be in G2/M phase, our results are consistent with the notion that the transcriptome of arrested cells may not necessarily mimic that of cycling cells (52, 53).

**Fig 5.**
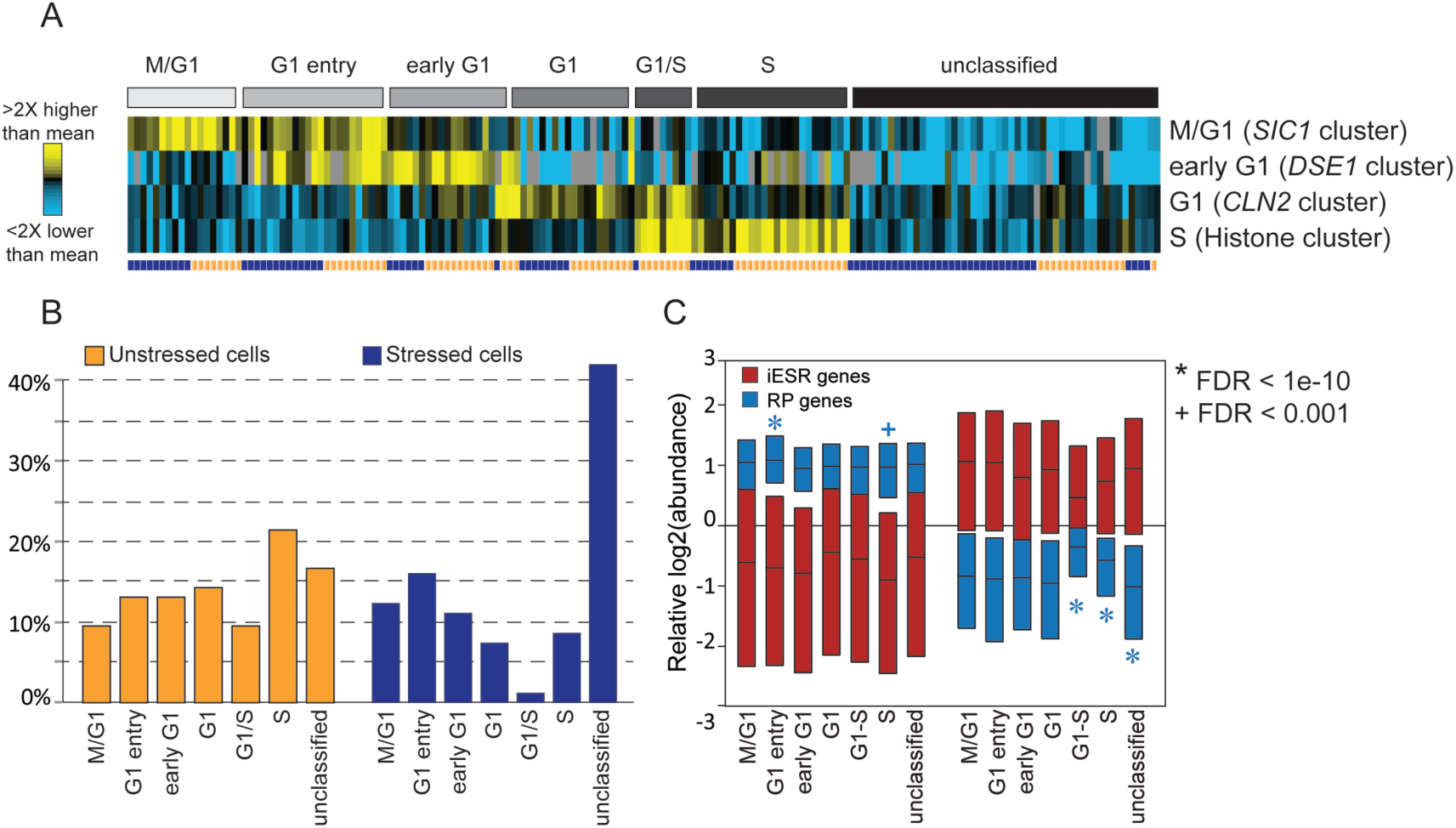
The influence of cell-cycle phase on ESR activation. Cycling genes used for classification were identified by clustering the scRNA-seq data (49) and then selecting clusters enriched for cell cycle markers (Table 3, see Methods). (A) Cells (columns) were clustered based on the centroid expression pattern of genes within each group (rows), and manually classified into and sorted within designated groups (A, grey bins). Stressed and unstressed cells are annotated by the purple/orange vector (A, bottom row). (B) The percentage of cells in each cell-cycle phase. (C) Boxplots (without whiskers) of all iESR (red) or RP (blue) genes from cells in that phase. Significance was assessed by Welch’s T-test on the pooled RP or iESR genes from cells within a given phase compared to all other cells; unstressed and stressed cells were analyzed separately. Note only one cell was classified as G1/S after stress.

We found no evidence that ESR activation as a whole is a function of the cell cycle. There were no statistically significant differences coordinately affecting iESR, RP, and RiBi transcripts at different points in the division cycle, including G1 phase. The only trend across the collective ESR groups was seen in stressed cells progressing through S phase at the time of collection: these cells showed significantly higher abundance of RP transcripts (p <1e-10) and, although not significant, slightly higher abundance of RiBi mRNAs and lower abundance of iESR transcripts (Fig S2). The simplest explanation is that these cells have partly acclimated and are thus relaxing ESR activation as they re-enter the cell cycle after G1 delay triggered by NaCl treatment (50, 51).

Although ESR activation as a whole was not coupled to cell cycle phase in unstressed cells, we were surprised to find concerted differences in RP expression, separable from activation of the ESR. Cells in early G1 had slightly, but statistically significantly, higher expression of RP transcripts; this correlates to the phase of maximal cell growth in yeast. In contrast, a third of unstressed cells in S-phase displayed concertedly low expression of RP mRNAs (FDR < 0.05, Fig S2) – these accounted for many of the cells identified in Fig 1C. The reduced expression of RP mRNAs was not related to higher iESR abundance or lower RiBi mRNA levels, aside of three cells in which the ESR appeared to be weakly activated (Fig S1). Thus, RP expression can be decoupled from ESR activation in a subset of unstressed cells.

### No evidence for mRNA cycling in the yeast metabolic cycle program

One potential link between RP expression and S-phase is the ultradian yeast metabolic cycle (YMC), which can be synchronized in bulk cultures through nutrient deprivation and has recently been reported in asynchronous, nutrient-replete cultures (54-57). In starvation-synchronized cultures, bulk transcriptome analysis identified three YMC phases, including an oxidative phase in which RP mRNAs peak, a reductive building phase in which respiration factors peak, and a reductive charging phase in which transcripts involved in fatty-acid metabolism and glycolysis are maximal (54). The reductive building phase is at least partly aligned with S-phase of the cell cycle (54, 55, 57, 58), which could explain why a subset of S-phase cells display low RP expression.

However, we did not find evidence for the same YMC transcriptome program reported in nutrient-restricted chemostats. First, there was no evidence that RP transcripts are cycling in our dataset. We used the program Oscope (59) to identify cycling transcripts, which were heavily enriched for cell cycle-regulated mRNAs (p = 2e-16, hypergeometric test (60)) but not RPs or transcripts encoding metabolic enzymes (Table 4). Second, we sought other RNAs whose patterns varied in accordance with RPs. Unstressed cells were ordered based on five representative RP transcripts using the WaveCrest algorithm (61), which then identified other mRNAs whose profiles fluctuated according to the same cell ordering (but not necessarily the same abundance profile, see Methods). Out of the top 100-ranked transcripts, most were RPs or mRNAs encoding translation factors (Table 5); one (*ENO1*) encoded a glycolysis enzyme and several were localized to mitochondria, but there was no enrichment for these categories. Finally, we looked explicitly at the relative abundances of RP, glycolysis, and other YMC mRNAs (Fig S3). There was cell-to-cell variation in abundance of glycolytic mRNAs, consistent with the YMC expectation; however, there was no statistically significant link to RP abundance. Furthermore, there was no evidence that other transcripts associated with the YMC oscillated in our study, either in abundance or detection rate within cells (Fig S3). Together, our results suggest that cells growing in rich medium may not display the same type of YMC-related transcriptome program as seen clearly in slow-growing nutrient restricted cells (54, 55).

### Heterogeneity in transcription factor targets implicates variation in regulation

We next searched for evidence of other cellular states that might influence expression of ESR gene groups. We leveraged the extensive knowledge of *S. cerevisiae* transcription factor (TF) targets to explore regulatory variation in single cells, by identifying cells with concerted expression differences in sets of TF targets, in two ways. First, we applied a gene-set enrichment approach to identify TF targets enriched among the distribution tails of relative log_2_ abundances in each cell (see Methods). Second, we applied a T-test per cell, comparing relative abundance of each group of TF targets compared to relative abundance of all other transcripts in that cell – although the latter approach may lack statistical power, it is sufficient to detect strong skews in TF-target behavior.

These approaches identified eight sets of TFs whose targets were coherently differently expressed in at least 3% of cells (Table 6). Several TF targets were differentially expressed in a large fraction of cells (Fig 6), including those of RP regulators (Ifh1, Fhl1, Rap1, Sfp1 (62, 63)), Dot6/Tod6 that repress a subset of RiBi genes during stress (64), and stress-responsive activators (Msn2, Hot1, Sko1 that regulate an overlapping set of targets). Several other factors were only implicated a subset of cells, including cell-cycle regulators as expected but also proteasome regulator Rpn4 and heat shock transcription factor Hsf1. In bulk RNA-seq experiments, proteasome genes appear weakly induced by NaCl (Fig S4) – our results instead show that Rpn4 is much more strongly activated in 11% of cells (FDR < 0.05), an effect that is lost in culture-level analysis. Hsf1 is not known to be activated by NaCl and its targets are not coherently induced in bulk RNA-seq experiments (Fig S4). Yet we identified higher expression of Hsf1 targets in 8% of stressed cells (FDR < 0.053), independent of Rpn4 target abundance. Thus, cells experience variation in signals related to protein degradation and folding in response to NaCl.

**Figure 6.**
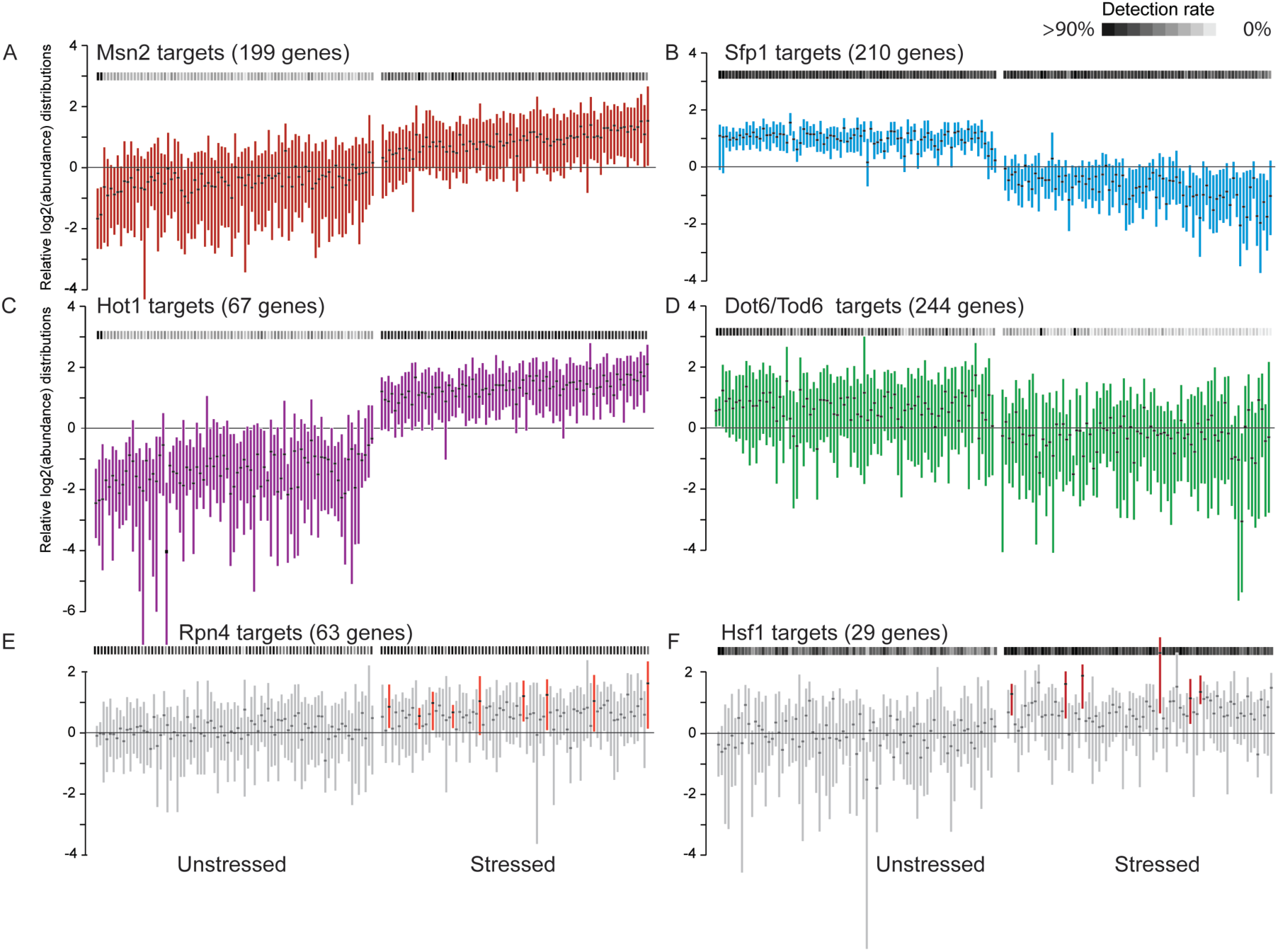
Regulatory variation across single cells. Distribution (without whiskers) of mean-centered log_2_(read count) values for indicated TF targets in single cells, organized as in Fig 1C. The number of targets for each TF is shown in parentheses. Gray-scale heat map (horizontal boxes) represents the detection rate, according to the key. (A-D) Targets of TFs that were differentially expressed in a large fraction of cells (see Table 6). (E-D) Cells for which targets of Rpn4 (E) or Hsf1 (F) were significantly elevated (FDR <0.053) compared to all other stressed cells are colored.

### Variation in TF relocalization reveals intrinsic and extrinsic variation in ESR regulation

To investigate the regulatory underpinnings of ESR variation revealed by scRNA-seq, we used single-cell microscopy to trace activation of the regulators implicated above. Cytosolic Msn2 and Msn4 rapidly relocalize to the nucleus upon various stress treatments; the same is likely true for Dot6 and Tod6 (26, 27, 65-67). In contrast, the rESR activator Sfp1 is nuclear during active growth, but ejected from the nucleus (and in some cases degraded) during stress to decrease RP transcription (24, 25, 68). Several upstream regulators also change localization during stress, notably the NaCl-activated Hog1 MAP kinase (69). While single-cell variation in Msn2/4 and Hog1 relocalization have been individually quantified (70-75), whether nucleo-cytoplasmic shuttling of these regulators before stress is coupled or fluctuates independently due to stochastic noise is not known. Furthermore, heterogeneity and dynamics of Dot6 and Sfp1 have not been investigated.

We therefore followed Msn2-mCherry localization in cells that also expressed Hog1, Dot6, or Sfp1 fused to GFP. We first quantified dual-factor localization in fixed cells. No cells showed nuclear Hog1 before stress, but 12% and 10% showed nuclear Msn2 or Dot6, respectively (based on identifiable nuclear objects, see Methods), consistent with the stressed state. However, only a third of cells with one factor localized to the nucleus also showed nuclear localization of the other. A small fraction (∼4%) of unstressed cells showed a dearth of nuclear Sfp1 signal (below the median ratio seen 30 min after stress, Fig 7A) – but there was no evidence of nuclear Msn2 in any of these cells. Upon NaCl treatment, the factors showed distinct dynamic behavior, with nuclear Hog1 peaking at 5 min, followed by maximal Msn2 and Dot6 nuclear localization at 15 min and 25 min, respectively (Fig 7B). Relocalization of Sfp1 was significantly prolonged and had not plateaued by 30 min after stress, consistent with the timing of transient rESR transcript reduction which troughs at 30-45 min after NaCl treatment (28).

**Fig 7.**
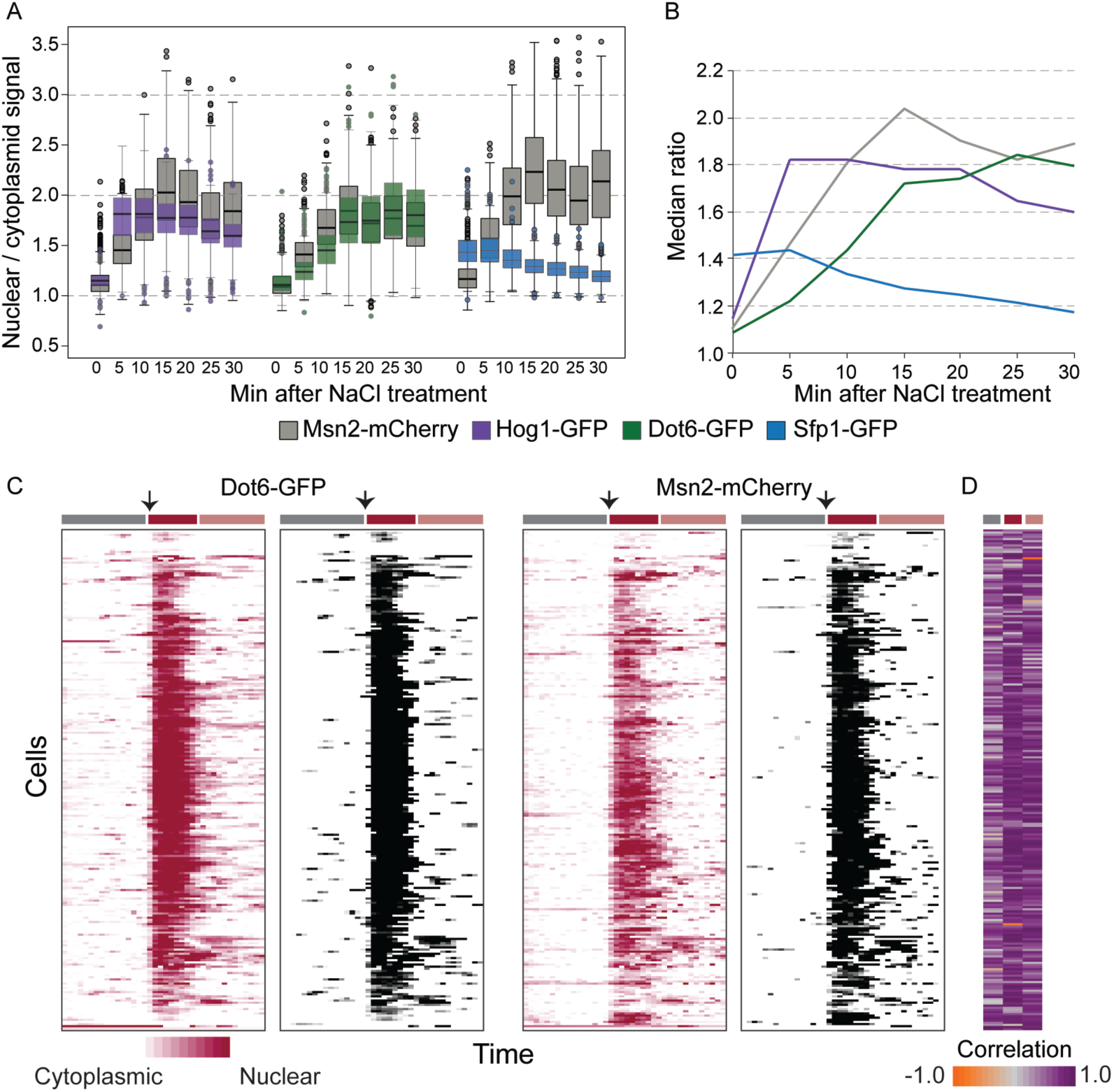
Stress-activated regulators show both coordinated and decoupled nuclear localization. (A) Distribution of nuclear/cytoplasmic signal for paired factors in individual cells before and after NaCl treatment (average n= 676 cells per time point). Data from two biological replicates were very similar and combined. (B) Median ratios from (A) plotted over time; Msn2 plot combines measurements from all three strains. (C) Nuclear TF signals (see Methods) of Dot6-GFP (left) and Msn2-mCherry (right) expressed in the same cells over time, before stress and after NaCl addition at 81 min (arrows). Each row aligned across all plots represents a different cell and each column represents a different time point. Red plots show traces of nuclear localization according to the key (see Methods), and corresponding grey-scale plots show quantitative measurements only for time points called as peaks. Colored boxes above the plots indicate 80 min before stress (grey box), 30 min after NaCl treatment (dark red box), and beyond 30 min after NaCl treatment (pink box). D) Correlation between Dot6-GFP and Msn2-mCherry traces for each temporal phase, according to the key.

We were especially interested in potential decoupling of ESR TFs, particularly in unstressed cells. However, differences in relocalization dynamics confound the analysis, since it could mimic decoupling in single-timepoint snapshots. We therefore followed TF dynamics in living cells, quantifying TF localization (see Methods) every 300 sec (to minimize light-induced stress (70)) and calling temporal peaks or troughs in nuclear concentration (see Methods). We were unable to confidently call troughs of nuclear Sfp1 before stress, but there appeared to be cells in which Sfp1 was depleted from the nucleus with no sign of nuclear Msn2 accumulation during the 80 min unstressed time course (Supplemental Movie 1). 19% and 22% of cells showed a detectible peak of nuclear Dot6-GFP or Msn2-mCherry, respectively, during 80 min of unstressed growth; but only 8% of cells showed nuclear translocation of both factors at some point during the experiment, generally with similar timing (median correlation in traces = 0.55, Fig 7C,D, Supplemental Movie 2). This fraction is higher than the joint probability of independent regulation (4%), and in close agreement with the 4% of cells for which scRNA-seq implicated weak ESR activation. Nonetheless, there were clear cases of decoupling: over a third of cells with a Dot6-GFP nuclear transition showed no called peak in nuclear Msn2-mCherry and low correlation (<0.2) in unstressed traces. These cells do carry unmarked Msn4, but it rarely transits to the nucleus in the absence of added stress (McClean, unpublished) and upon stress treatments generally correlates closely with Msn2 (67). In all cases studied here, pre-stress nuclear pulses were both shorter and milder quantitatively compared to after NaCl treatment. Both factors transited to the nucleus upon NaCl treatment in essentially all cells, after which time cells showed nucleo-cytoplasmic bursts that were partly decoupled (across cells and factors), as previously reported for Msn2 (76). Thus, while Dot6 and Msn2 activation were highly correlated during the acute response to NaCl, the pre-stress and post-acclimation phases showed evidence of both coordinated and decoupled nuclear fluctuations of the regulators.

## DISCUSSION

Our work addresses several unanswered questions regarding heterogeneity in stress defense and tolerance. Many past studies have characterized the transcriptomic responses to stress at the culture level, presenting a wealth of information on the mode and mechanisms of stress defense. A critical missing component from past bulk studies is how and why individual cells vary in their response. Our results indicate that individual yeast cells can vary substantially in the magnitude of their transcriptome response, both before and after stress, and that individual cells experience stress differently (exemplified by quantitative differences in ESR activation and differential expression of proteasome and chaperone-encoding transcripts after NaCl treatment). The extensive knowledge of yeast transcriptional regulation enabled us to investigate sources of upstream transcriptome regulation, implicating heterogeneity both intrinsic to individual regulatory paths and extrinsic to the cellular system.

### Heterogeneous ESR activation before stress likely influences stress survival

Both the scRNA-seq results and fluorescent-TF profiling suggest that a subset of cells mediate mild activation of the ESR in the absence of added stress. Four percent of cells showed mildly higher iESR and lower RP mRNA abundance (Fig 1C), consistent with the 4% of cells estimated to co-regulate iESR/rESR regulators Msn2 and Dot6 (Fig 7C). We propose that this mild activation underlies the heterogeneity in single cell survival of extreme stress doses. Although we observed decoupled Msn2 and Dot6 pre-stress nuclear fluctuations by microscopy, we did not observe decoupled activation of their combined targets in individual unstressed cells (Fig 1 and unpublished). The amplitude and duration of pre-stress Msn2/Dot6 pulses was significantly smaller and shorter than after NaCl stress. In the case of Msn2, relocalization is influenced by both nuclear import and export that together produce distinct temporal profiles (71, 77-79). One prediction is that genes with more Msn2 binding sites are more sensitive to brief pulses of nuclear Msn2 (20, 80, 81). However, our data did not support this: genes with many Msn2 binding sites showed no more evidence of concerted pre-stress fluctuations than genes with few binding sites (Fig S5). Thus, cells likely maintain a filtering system to distinguish a true upstream signal from noisy TF activation. This system could emerge from chromatin regulation (80, 82) or other regulatory signals (*e.g.* posttranslational TF modification) that act as gate keepers to the transcriptome response.

A remaining question is why some unstressed cells activate the ESR program. One model is that stochastic fluctuations in a common upstream regulator produce stochastic but coordinated activation of the downstream factors. A candidate is Protein Kinase A, which phosphorylates and suppresses several stress-activated regulators (including Msn2 and Dot6), promotes expression of RP transcription (83, 84), and has been implicated in stochastic Msn2 regulation (65, 79, 85, 86). Whether PKA fluctuations represent random events or respond to some cellular signal is not clear. A second, compatible model is that cells with mild ESR activation are actually experiencing, and thus actively responding to, internal stress. Such stress could emerge from normal cellular processes, *e.g.* damage from bursts of oxidative metabolism or during DNA replication. A third model that our data discounts is that the ESR fluctuates with the cell cycle in normally dividing cells (35). We did see a milder ESR activation in post-stress cells in S-phase, but we believe this is merely a sign of acclimation-dependent re-entry of the cell cycle. We propose that the previously reported correlation between ESR activation and prolonged G1 in mutants is likely a response to deleterious gene deletions rather than an inherent coupling of the ESR to G1 phase.

### Exquisite control of RP transcripts can be decoupled from the ESR

In many bulk transcriptomic yeast studies to date, RP expression is inversely proportional to stress-defense transcripts in the iESR, and these gene groups display opposing responses during rapid growth *versus* environmental stress (21, 34, 36). Indeed, these gene groups are controlled by the same upstream signaling pathways, including PKA, TOR and stress-activated regulators (87-89). But studying individual cells expands knowledge of the regulatory system: although RP and iESR transcripts are anti-correlated in most cells in our analysis, RP expression is clearly decoupled in a subset of unstressed individuals. The reason and mechanisms remain unclear. The link between low RP transcripts and S-phase is appealing: although we found no clear evidence for the same YMC transcriptome seen in nutrient-synchronized cultures (54), metabolic genes are regulated during the G1/S transition (90, 91). The yeast CDK, Cdc28, also binds to RP promoters (92), and an imbalance of ribosome components can trigger G1/S delay (93, 94). Future work will be required to decipher this regulation, as well as the mechanisms that give rise to exquisite control minimizing variation and ensuring cell presence of RP mRNAs.

### Implications for heterogeneous stress responses in other organisms

Heterogeneity in microbial stress tolerance has been proposed to serve as a bet-hedging mechanism, ensuring that a minimal fraction of the population survives in the event of catastrophic environmental events (6, 7, 10). But the phenomenon is also observed in multi-celled mammalian systems (2-4, 95) and at least partly influenced by variable activation of the p53 tumor suppressor. Like Msn2, inactive p53 resides in the cytoplasm but upon stress rapidly relocates to the nucleus with transient pulses, where it activates target-gene transcription and has been reported to repress ribosome-producing polymerase I and III (89, 96-98). p53 targets with multiple and high-affinity binding sites are most sensitive to transient nuclear bursts, as seen for Msn2 targets, whereas other genes require prolonged p53 activation for full induction (96, 99, 100). And, as is the case with Msn2 activation, prior induction of p53 leads to subsequent tolerance to what would otherwise be lethal drug doses (101). P53 also shows heterogeneous nuclear pulses in proliferating cells without exogenous stress. Unlike Msn2, which shows quantitatively shorter and weaker pre-stress nuclear pulses, the amplitude and duration of pre-stress p53 pulses is similar to that seen after inflicted stress; but like the yeast factor these pre-stress bursts do not necessarily alter gene expression (102). Instead, layers of post-translational p53 modification can filter potential noise in the regulatory system. A better understanding of the regulatory networks that control heterogeneity in transcription and stress tolerance is likely to open new avenues to control population behavior.

## MATERIALS AND METHODS

### Strains and growth conditions

All experiments were done in the BY4741 background. Unless noted, cells were grown in rich YPD medium in batch culture at 30C for at least 7 generations to mid-log phase, at which point an aliquot was removed to serve as the unstressed sample. NaCl was added to 0.7M to the remaining culture and cells were grown for 30 min. Unless otherwise noted for specific applications, cells were collected by brief centrifugation, decanted, and flash frozen in liquid nitrogen. Strains expressing tagged proteins were generated by integrating an mCherry-*HIS3* cassette downstream of *MSN2* in BY4741 strains from the GFP-tagged collection (103), which were verified to harbor the GFP-*HIS3* cassette downstream of *DOT6*, *SFP1*, or *HOG1* (generating strains AGY1328, AGY1329, and AGY1331, respectively).

### Single-cell sorting, library preparation and sequencing

Fluidigm’s C1 microfluidic platform was adapted to perform cDNA synthesis from single yeast cells. Flash frozen cells were re-suspended in 1 mL of 1 M Sorbitol on ice, counted on a hemocytometer, and then diluted to approximately 4x10^5^ cells per mL in a final volume of 200 μL. To generate partial spheroplasts that can easily lyse on the Fluidigm C1 microfluidic device, we titrated each sample with different amounts of zymolyase (0.025U, 0.0125U, 0.00625U, and 0.003125U) and incubated cells for 30 min at 37°C. This was done because unstressed and stressed cells displayed different sensitivities to zymolyase digest. After incubation, cells were spun at 250 g for 4 min and re-suspended in Sorbitol Wash buffer (0.455x C1 Cell Wash Buffer, 1M sorbitol, 0.2ug/μl BSA, 0.08(8)U/μL SUPERase RNAse Inhibitor). Samples with the maximal number of intact spheroplasts (compared on a Leica DMI 6000 inverted microscope) were diluted to a final concentration of 600 cells/μL, and 9 μL of these cells were mixed with Fluidigm Suspension reagent at final loading concentration of 275 cells/μL and loaded onto the primed C1 Chip designed to capture 5-10 μm cells, according to manufacturer instructions. The cell concentration in the loading mixture was crucial to maximize the number of wells capturing single cells inside the microfluidic device. Another modification was that 1M Sorbitol was added to all wash buffers to prevent premature lysis. After cell loading, each chip was visually inspected and imaged to tabulate single-cell capture rates. Roughly 50% of wells contained a single cell, verified by imaging and manual inspection. This rate is lower than the normal capture rate because yeast cells are smaller and deform less. Spheroplasts were lysed in the Fluidigm instrument and cDNA was generated using Clontech reagents for Fluidigm C1 based on the single-cell RNA-seq protocol (cat # 635025). Finally, cDNA was harvested from each Fluidigm C1 chip and into a 96 well plate for storage at −20C. ERCC spike-in sequences (mix A) were added at 1:4x10^5^/μL of the concentration provided in the original product (Ambion catalog number 4456740).

Before library preparation, cDNA from each cell was quantified on an AATI Fragment Analyzer. Using concentrations calculated from a smear analysis between 450bp and 4500bp, cDNA from each cell was diluted with TE to ∼0.2 ng/μL using the Mosquito X1 pipetting robot (TTP Labtech). Diluted cDNA served as the template for Nextera XT library generation following manufacturer protocol (Illumina catalog number FC-131-1096) with some modifications. Because we used TTP’s Mosquito HTS 16 channel pipetting robot (capable of accurately aliquoting volumes down to 50nL), we were able to scale down the total volume of each Nextera XT library to 4μL. More specifically, for each 400 nL of input template DNA, we added 400 nL Tagmentation mix and 800 nL Tagmentation buffer in a final volume of 1.6 nL. The Tagmentation reaction was incubated at 55°C for 10 min. Neutralization was done by adding 400 nL Neutralization buffer to the above reaction and incubating 10 min at room temperature followed by the addition of primers at 400 nL each and NPM PCR master mix at 1200 nL. The PCR step was run for 10 cycles. 1 μL of each library was combined to form two separate pools, one for unstressed cells and one for stressed cells. Two rounds of size selection was performed using Agencourt AMPure beads (Beckman Coulter catalog number A63882). 100 ng of each pool was combined and sequenced in one run on 3 lanes of an Illumina HiSeq-2500 1T v4 sequencer for 150-bp paired-end sequencing. Reads generated across the three lanes were merged and de-multiplexed using Illumina software bcl2fastq v1.8.4, allowing no mismatches and excluding the last position (8^th^ index base).

Paired-end reads were mapped to the S288c *Saccharomyces cerevisiae* genome R64-2-1 (39, 104) with ERCC spike-in sequences added using BWA mem Version: 0.7.12-r1039 and default parameters (104). Reads were processed with Picard tools Version: 1.98(1547) cleansam and AddOrReplaceReadGroups as required by downstream applications. Resulting bam files were sorted and indexed using Samtools Version 1.2. Paired-end fragments were deduplicated using RemoveDuplicate function in Picardtools, and read counts mapped to genes were extracted using FeatureCounts Version 1.5.0. Sequenced wells were removed from the analysis if they had <1,000 total mapped reads or if the proportion of ERCC spike-ins to total-mapped reads was >0.2 (105). Data were normalized by SCNorm (106) in R version 3.3.1; ERCC spike-in samples were not used in the normalization. Normalized read counts for each gene were logged and then centered by subtracting the mean log_2_(read counts) for that gene across all cells in the analysis. All data are available under GEO access number GSE102475.

### Statistical analysis of differential expression

Individual cells with altered expression of defined gene groups (*e.g.* cells with low RP expression as in Fig 1C or high Rpn4/Hsf1 targets in Fig 6) were determined using a two-tailed Welch’s T-test, comparing the set of mean-centered log_2_(read count) values in that cell to the combined set of mean-centered log_2_(read count) values for all other unstressed (or stressed) cells, taking FDR <5% (107) as significant. To score differential expression of ESR groups across cell-cycle phases, a two-tailed Welch’s T-test was applied to the mean-centered log_2_(read count) values, comparing the pooled set of values from all cells in a given cell-cycle phase to the pooled values from unstressed or stressed cells from all other phases; stressed and unstressed cells were analyzed separately unless otherwise noted. All cell classifications from this work are summarized in Table 8.

The Oscope (59) R package version 1.4.0 was used to identify oscillatory genes in the set of unstressed cells. Oscope first filtered transcripts using the function CalcMV, and analyzed only those having a minimum mean larger than 15 (MeanCutLow=15 and otherwise default parameter settings). Oscope then fit sinusoidal functions to all remaining mRNA pairs and those identified as oscillating were clustered according to their oscillation frequencies. Oscope was also run with relaxed mean and variance thresholds to consider all genes with a mean larger than 10 and the maximum number of clusters set to five in the K-medioids clustering step (Table 4). Oscope computationally reordered the single-cells for the two detected gene clusters. The cyclic orderings were used to identify additional genes following the same orders by fitting a 3^rd^ degree polynomial to all genes using the WaveCrestIden function from the WaveCrest (61) R package version 0.0.1. Genes were ranked by their fit using the mean squared error (MSE) and only the top 100 genes were considered further. The WaveCrestIden function was run twice, either including zeros in the polynomial fit or treating them as missing. The WaveCrest algorithm was also used as above to obtain a cyclic order on the set of unstressed single-cells based on five ribosomal protein (RP) genes (Table 5).

### Variance and detection-rate analyses

Length-normalized read counts were taken as SCNorm-normalized read counts per transcript divided by transcript length, and the mean (or median) for each transcript across all unstressed or stressed cells was calculated; mean and median values were essentially the same (R^2^ = 0.99). Unless otherwise noted, transcripts with no read count in that sample were not included in the calculation (instead of counting the value as 0) – the only exception was in calculating correlations between average scRNA-seq data compared to bulk data (*e.g.* Fig S4), for which the correlation was significantly higher by scoring no-read transcripts as zero values and including them in the calculation. Detection rate was defined as the fraction of unstressed or stressed cells in which a gene was detected by at least one collapsed read count. We devised a statistic to test if RP, RiBi, or iESR gene groups were significantly different from other genes. For each set of genes, a cubic smoothing spline was fit to describe the relationship between detection rate and median expression, and the point along the curve at which 80% of points were fit was identified. This process was repeated for 10,000 random gene sets equal to the size of the query gene group. The p-value was calculated as the fraction of random gene sets having a statistic more extreme than the observed value. The calculated statistic for unstressed and stressed cells, respectively was: RP: 0.045, 0.089; RiBi: 0.148, 0.156; iESR: 0.493, 0.208. RP genes were also compared against 10,000 trials randomly selecting mRNAs shorter than the median-RP gene length. Tests were repeated on genes without a close homolog in the S288c genome (*i.e.* genes with BLAST hits of E>1e-5). All significant tests shown in Fig 3 remained significant (p<0.05) except for RP transcripts after stress (p = 0.36).

### Cell classifications and gene clustering

Data were clustered using Pagoda (49) using default parameters and clusters enriched for known cell-cycle regulators (60) or glycolysis transcripts were manually identified. We were unable to identify known cell cycle markers that peaked in M phase, either in the Pagoda-clustered data or using known M-phase markers (60). To classify cells according to cell cycle phase, cells were organized by hierarchical clustering (108) based on the centroid (median) of mean-centered log_2_(transcript abundance) for transcripts in each cell-cycle phase (Table 3) and manually sorted and classified based on overlapping peaks of each vector (Fig 5A). Cells that showed no expression peak in any of the vectors were scored as ‘unclassified’.

### Identification of TF target expression differences

Compiled lists of TF targets were taken from (88). We added to this an additional list of Msn2 targets, taken as genes with stress-dependent Msn2 binding within the 800 bp upstream region and whose normal induction during peroxide stress required *MSN2/MSN4* ((109), Table 7) and genes whose repression requires the *DOT6/TOD6* repressors (defined here as genes with a 1.5X repression defect in two biological replicates of wild-type BY4741 and *dot6Δtod6Δ* cells responding to 0.7M NaCl for 30 min ((110), Table 7). In total, we scored 623 overlapping sets of TF targets defined in various datasets (109, 111-113) or summarized from published ChIP studies (114). We identified TF targets with concerted expression changes in two ways. First, we identified the distribution tails in each cell, identifying all mRNAs in that cell whose mean-centered log_2_(read count) was ≥1.0 or ≥2.0 (*i.e.* 2X or 4X higher than the population mean). We then used the hypergeometric test to identify sets of TF targets enriched on either list, taking the lower of the two p-values for that TF-cell comparison. Comparable tests were done for the sets of genes whose relative abundance was ≤-1.0 or ≤-2.0 in each cell. TFs with -log_10_(p-value)>4 were taken as significant (equivalent to a cell-based Bonferroni correction, p = 0.05 / 623 tests = ∼1e-4). We focused on TFs whose targets were enriched at the distribution tails in at least 4 cells, which is unlikely to occur by chance). Two sets of TF targets were removed because their targets heavily overlapped with ESR targets and their enrichment was not significant when those overlapping mRNAs were removed (Table 6). In a second approach, we used Welch’s T-test to compare the mean-centered log_2_(read count) values of each set of TF targets compared to all other measured mRNAs in that cell, again taking -log_10_(p-value)>4 as significant and focusing on TFs identified in at least 3 cells. Finally, for Fig 6E ad 6F colored boxplots indicate TF targets whose relative abundances in the denoted cell were significantly different from all other stressed cells (FDR < 0.053).

### FISH

BY4741 was grown as described above and collected for fixation and processing as previously described (115). FISH probes were designed using the Biosearch Technologies Stellaris Designer with either Quasar 670, CAL Fluor Red 610, or Quasar 570 dye, using 33 probes for *PPT1* (Quasar 670) and *RLP7* (Quasar 570) and 48 probes for *SES1* (Quasar 570). *PPT1* and *RLP7* were measured in the same cells, and *SES1* was measured in a separate experiment as a control. Images were acquired with as z-stacks every 0.2 mm with an epifluorescent Nikon Eclipse-TI inverted microscope using a 100x Nikon Plan Apo oil immersion objective and Clara CCD camera (Andor DR328G, South Windsor, CT. Quasar 670 emission was visualized at 700 nm upon excitation at 620 nm (Chroma 49006_Nikon ET-Cy5 filter cube, Chroma Technologies, Bellows Falls, VT, USA). Quasar 570 emission was visualized at 605 nm upon excitation at 545 nm (Chroma 49004_Nikon ET-Cy3 filter cube). *PPT1* and *RLP7* transcripts were counted manually, while *SES1* mRNA was counted by semi-automated transcript detection and counting in MATLAB using scripts adapted from (116).

### Fixed-cell microscopy

Cells were grown as described above and fixed with 3.7% formaldehyde for 15 min either before or at indicated times after NaCl addition. Cells were washed 2X with 0.1M potassium phosphate buffer pH 7.5, stained 5 min with 1 μg/mL DAPI (Thermo Scientific Pierce, 62247) and additionally washed 2X with 0.1M potassium phosphate buffer pH 7.5. Cells were imaged on an epifluorescent Nikon Eclipse-TI inverted microscope using a 100x Nikon Plan Apo oil immersion objective. GFP emission was visualized at 535 nm upon excitation at 470 nm (Chroma 49002_Nikon ETGFP filter cube, Chroma Technologies, Bellows Falls, VT, USA). mCherry emission was visualized at 620 nm upon excitation at 545 nm (Chroma 96364_Nikon Et-DSRed filter cube). DAPI emission was visualized at 460 nm upon excitation at 350 nm (Chroma 49000_Nikon ETDAPI filter cube). Nuclear to cytoplasmic intensity values were calculated with customized CellProfiler scripts (117). The fraction of cells with nuclear factor before stress was calculated by identifying nuclear masks in Cell Profiler (*i.e.* identifiable nuclear objects) in that channel that overlapped with DAPI masks and manually correcting miscalls.

### Live-cell microscopy

Yeast strains AGY1328 (Dot6-GFP Msn2-mCherry) and AGY1331 (Sfp1-GFP Msn2-mCherry) were grown at 30°C with shaking to OD_600_ 0.4-0.5 in low-fluorescence yeast media (115)(LFM); the fraction of cells with nuclear Msn2/Dot6/Sfp1 was very similar in fixed, unstressed cells growing in LFM versus YPD, not shown. Cells were imaged in a Focht Chamber System 2 (FCS2) (Bioptechs, Inc; Butler, PA) with temperature maintained at 30° C. Cells were loaded into the chamber by adhering them to a 40mm round glass coverslip. Briefly, the coverslip was prepped by incubating with concanavalin A solution (2mg/ml in water) for 2 minutes at room temperature. The concanavalin A was aspirated, and 350μl of cell culture added. Cells were allowed 2 minutes at room temperature to adhere to the concanavalin A before the gasket was washed once with 350μl of fresh LFM media. A round 0.2μM gasket was used to contain the concanavalin A solution and cell solution. The entire assembly, including coverslip, cells, and gasket was assembled with the microaqueduct slide, upper gasket, and locking base as per manufacturer instructions to create an incubated perfusion chamber. This assembly was transferred to an epifluorescent Nikon Eclipse-IT inverted microscope for time-lapse imaging. Temperature was maintained at 30°C using the Bioptechs temperature controller. Steady perfusion with LFM or LFM + 0.7M NaCl (initiated at 81 min) was maintained utilizing FCS2 micro-perfusion pumps feeding media through the FCS2 perfusion ports. Once loaded onto the microscope, cells and temperature were monitored for at least one hour to ensure stable temperature and robust growth.

During a timecourse experiment, a TI-S-ER motorized stage with encoders (Nikon MEC56100) and PerfectFocus system (Nikon Instruments, Melville, NY, USA) were used to monitor specific stage positions and maintain focus throughout the timecourse. A Clara CCD Camera (Andor DR328G, South Windsor, CT) was used for imaging. GFP emission was visualized at 535 nm (50 nm bandwidth) upon excitation at 470 nm (40 nm bandwidth; Chroma 49002_Nikon ETGFP filter cube, Chroma Technologies, Bellows Falls, VT, USA). mCherry emission was visualized at 620 nm (60 nm bandwidth) upon excitation at 545 nm (30 nm bandwidth; Chroma 6364_Nikon Et-DSRed filter cube). Single-cell traces of nuclear localization were extracted from fluorescence images using custom Fiji (118) and Matlab scripts. Fiji was used to threshold and identify individual cells. Individual cell tracks were constructed using a modified Matlab particle tracking algorithm (119) and manually validated and corrected. Localization of individual transcription factors was quantified using a previously published localization score consisting of the difference between the mean intensity of the top 5% of pixels in the cell and the mean intensity of the other 95% of pixels in the cell (120). Localization of transcription factors to the nucleus was identified using findpeaks2 (Matlab File Exchange) to identify peaks in the transcription factor localization score corresponding to nuclear localization of the transcription factor of interest.

## ACKNOWLEDGEMENTS

This work was supported by: R01GM083989 to APG, Department of Energy Great Lakes Bioenergy Research Center (DOE Office of Science BER DE-FC02-07ER64494), and by a Career Award at the Scientific Interface from the Burroughs Wellcome Fund to MNM; work conducted by the U.S. Department of Energy Joint Genome Institute, a DOE Office of Science User Facility, was supported by the Office of Science of the U.S. Department of Energy under Contract No. DE-AC02-05CH11231.

## AUTHOR CONTRIBUTIONS

APG designed and oversaw the project, performed downstream analysis, and wrote the manuscript, with comments from other authors. FBY and JK optimized and performed the Fluidigm capture and cDNA generation under the supervision of SRQ. DC and LS performed library generation and sequencing under the guidance of IG. JH generated cell samples for scRNA-seq, generated fluorescently tagged strains, and performed fixed-cell microscopy; LEE performed FISH experiments; MP performed scRNA-seq read mapping and clustering and developed and applied CellProfiler scripts; RB performed scRNA-seq normalization and statistical analysis under the supervision of CK. MNM designed and performed live-cell TF imaging and analysis and supervised FISH experimental design and application.

## FIGURE LEGENDS

**Figure S1.**
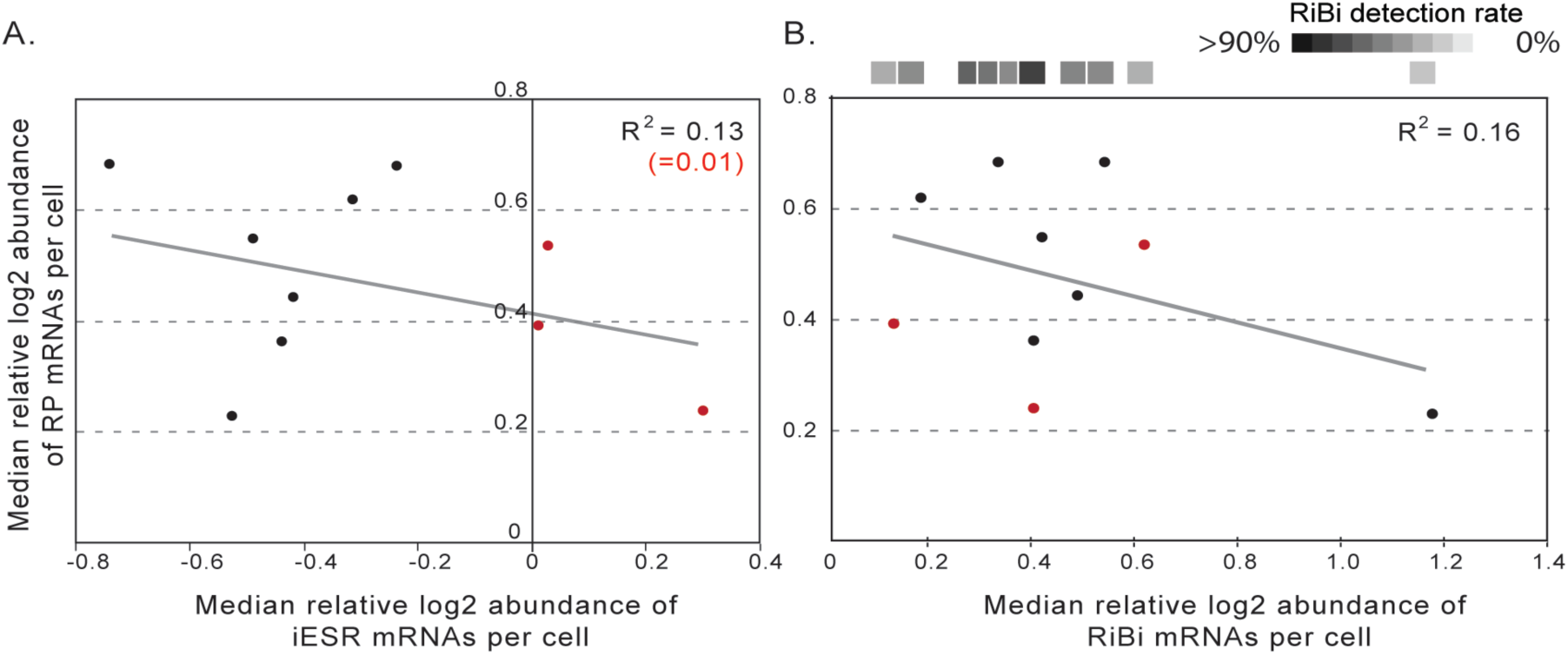
For each cell plotted, the median of the mean-centered log_2_(read count) values for the group of RP transcripts was plotted against the median of the mean-centered log_2_ (read count) values for (A) iESR transcripts and (B) RiBi transcripts in that cell. Red points represent the cells in which iESR transcripts were concertedly high (Fig 1C, asterisks). The RiBi-transcript detection rate (% of transcripts measured) for each cell is represented above the corresponding points in B, color-coded according to the key. R^2^ is shown for all points and excluding red points in which iESR transcripts were relatively high.

**Figure S2.**
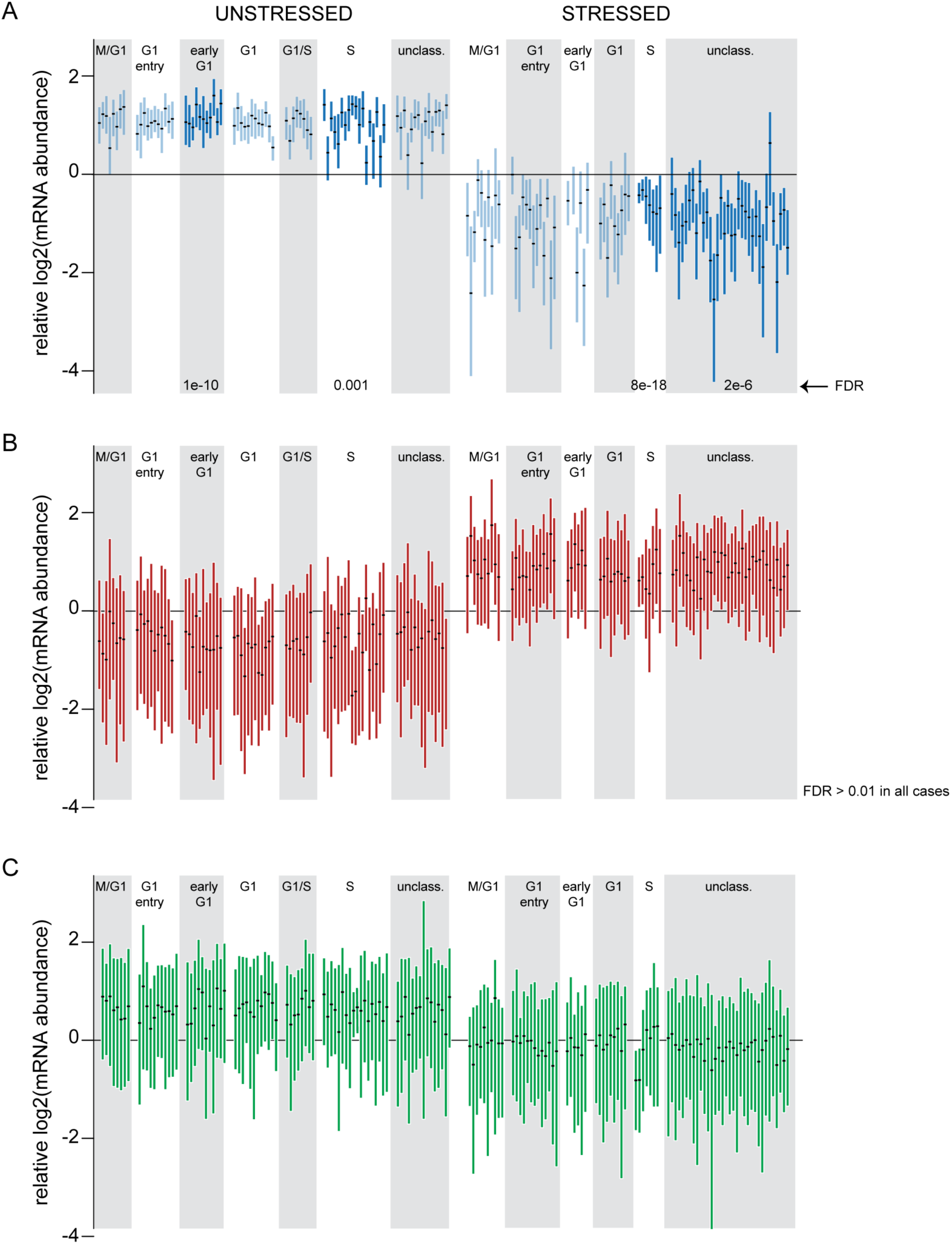
Boxplots (without whiskers) of (A) RP, (B) iESR, (C) RiBi mRNAs for cells classified in different cell cycle phases. Each boxplot represents the distribution of relative mRNA abundances, as defined in Methods, within each cell. Significance was assessed by Welch’s T-test on the pooled RP, iESR, or RiBi genes from cells within a given phase compared to the pooled set of those transcripts from all other cells; unstressed and stressed cells were analyzed separately. Dark blue boxplots in (A) represent significant groups, with FDR listed below. RP expression was slightly, but highly statistically significantly, higher among cells in G1 phase, particularly for subsets of cells. A subset of unstressed cells in S-phase showed particularly low mean-centered log_2_ abundance of RP transcripts. In stressed cells, those in S-phase had particularly tight distribution of RP abundance, showing in effect weaker RP repression during stress than cells in other phases. There were no significant differences across cell-cycle phases for expression of iESR or RiBi transcripts.

**Figure S3.**
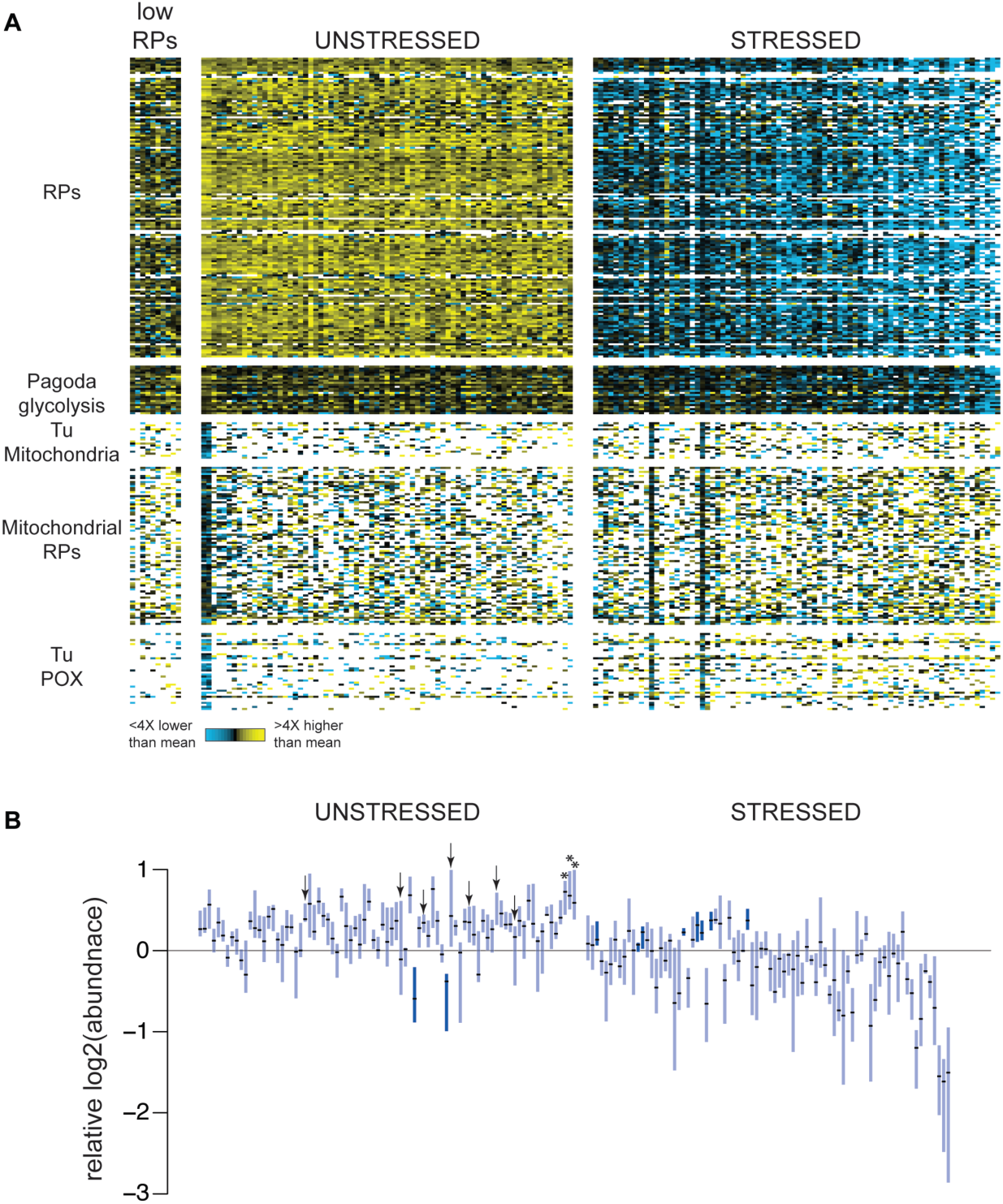
(A) Mean-centered log_2_(read count) values of several groups of transcripts, in unstressed cells with low RPs, other unstressed cells, and stressed cells, as described in Figure 1C. Cells are ordered as in Figure 1, except that unstressed, low-RP cells are displayed first. Gene groups include: 149 mRNAs annotated as part of the ESR RP cluster, 28 genes identified by Pagoda clustering that are heavily enriched for glycolysis transcripts, 23 and 40 YMC-related mRNAs identified by Tu *et al.* enriched for mitochondrial or peroxisomal (POX) functions; also shown are 82 transcripts encoding mitochondrial ribosomal proteins. (B) Boxplots (without whiskers) of the distribution of mean-centered log_2_(read counts) for the group of Pagoda-identified glycolysis transcripts, in individual cells ordered as in Figure 1C. Cells with statistically different expression of glycolysis mRNAs (FDR < 0.05) are highlighted in dark blue (unstressed and stressed cells were analyzed separately). Cells with lower RP expression from Fig 1C are indicated with arrows and asterisks, as described in Fig 1C. Cells with low-RP expression had no statistically significant difference in glycolysis-mRNA expression.

**Figure S4.**
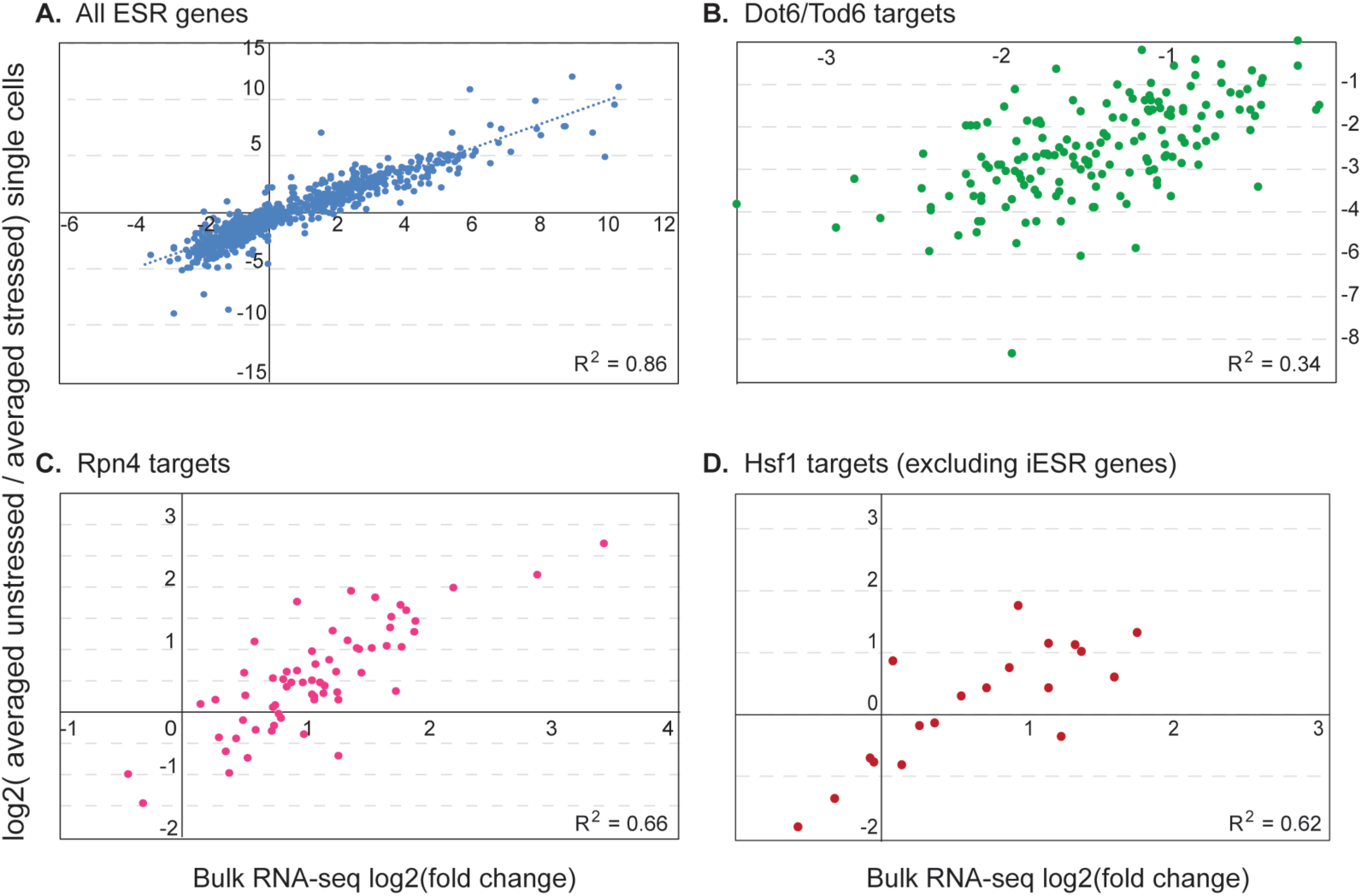
Correlation between bulk RNA-seq log_2_(fold-change) after 0.7M NaCl and average of single cell measurements. Bulk data represent the average of three biological replicates (Ho *et al*, 2017, submitted). R^2^ is shown for each plot. Correlations were substantially higher when transcripts with no reads in a given cell were scored as zero reads (rather than treated as missing values), which supports the notion that missing measurements are influenced biological variation in transcript presence per cell, rather than purely noise specific to scRNA-seq data.

**Figure S5.**
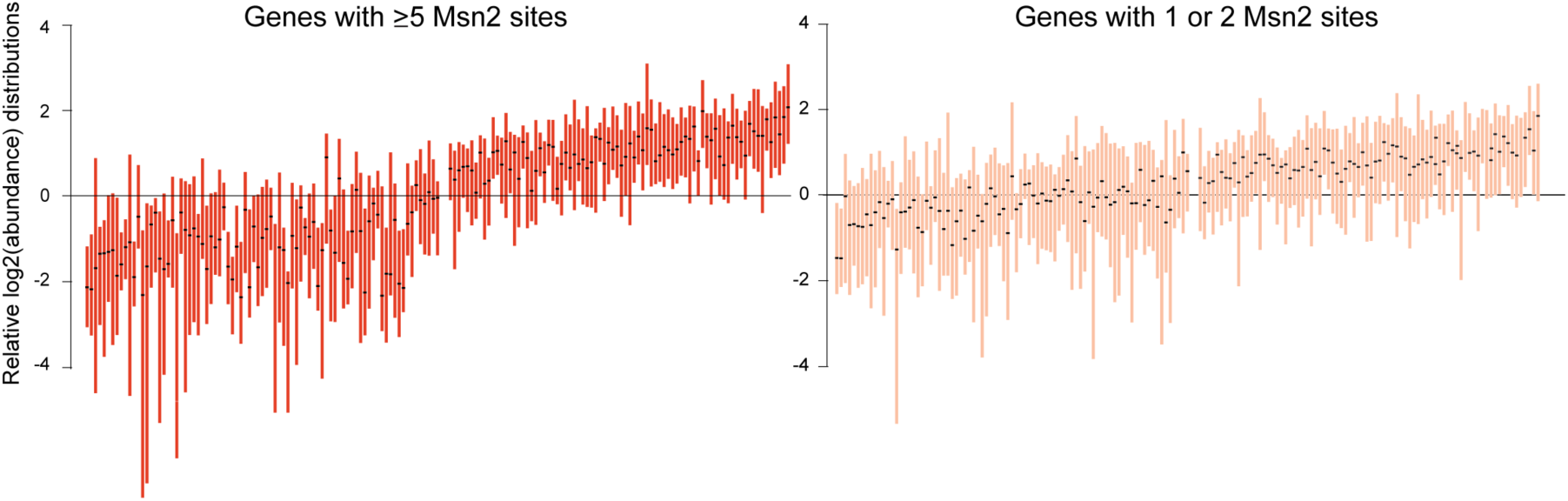
Genes with more upstream Msn2 sites do not show significant differences in pre-stress activation. The number of Msn2 binding sites (CCCCT or CCCCC) within 800bp upstream of the gene starts was identified in Msn2 targets defined by Huebert *et al* (2012). The distribution of mean-centered log_2_(read counts) in unstressed cells for genes with at least 5 upstream elements (left) was not statistically significantly different from genes with one or two upstream elements (right)(Welch’s T-test comparing the two groups in each cell). Cells are ordered as in Fig 1C. None of the cells showed concerted differences in Msn2 target expression before stress (FDR < 0.05, Welch’s T-Test, see Methods). There was no relationship between the median relative log_2_ read counts and cells with low RPs.

**Supplemental Movie 1**. Representative timecourse of Msn2-mCherry and Sfp1-GFP before and after addition of 0.7M NaCl. The pre-stress phase is indicated by a white scale bar, which turns red upon NaCl addition. We were unable to call troughs of nuclear Sfp1-GFP before stress. Apparent Sfp1-GFP nuclear depletion, without nuclear Msn2-mCherry during the unstressed phase, is seen in the cell at the far upper left corner; the nucleus is clearly in focus judged by the nuclear Msn2-mCherry concentration upon NaCl treatment.

**Supplemental Movie 2**. Representative timecourse of Msn2-mCherry and Dot6-GFP before and after addition of 0.7M NaCl. The pre-stress phase is indicated by a white scale bar, which turns red upon NaCl addition.

## List of Tables and Files

Table 1: Statistics on sequencing features

Table 2: Transcripts outside RP-splines and iESR-fit splines

Table 3: Cell-cycle expression vectors and gene lists used for classification

Table 4: Oscope-identified clusters of cycling genes

Table 5: Wavecrest-identified transcripts associated with RP mRNAs

Table 6: Specific TF-target gene sets identified in Fig 6 analysis

Table 7: Lists of all TF targets considered for Fig 5 analysis

Table 8: Summary of all cell classifications

Move 1: Representative Msn2-mCherry and Sfp1-GFP before and after addition of 0.7M NaCl

Movie 2: Representative Msn2-mCherry and Dot6-GFP before and after addition of 0.7M NaCl

